# Transcriptomics of developing wild sunflower seeds from the extreme ends of a latitudinal gradient differing in seed oil composition

**DOI:** 10.1101/2021.06.08.447625

**Authors:** Max H. Barnhart, Edward V. McAssey, Emily L. Dittmar, John M. Burke

## Abstract

Seed oil composition, an important agronomic trait in cultivated sunflower, varies latitudinally across the native range of its wild progenitor. This pattern is thought to be driven by selection for a higher proportion of saturated fatty acids in southern populations compared to northern populations, likely due to the different temperatures experienced during seed germination. To investigate whether these differences in fatty acid composition between northern and southern populations correspond to transcriptional variation in the expression of genes involved in fatty acid metabolism, we sequenced RNA from developing seeds of sunflowers from Texas, USA and Saskatchewan, Canada (the extreme ends of sunflower’s latitudinal range) grown in a common garden. Over 4,000 genes were found to be differentially expressed between Texas and Canada, including several genes involved in lipid metabolism. Many differentially expressed oil metabolism genes colocalized with known oil QTL. The genes producing stearoyl-ACP-desaturases (*SAD*) were of particular interest because of their known role in the conversion of fully saturated into unsaturated fatty acids. Two *SAD* genes were more highly expressed in seeds from Canadian populations, consistent with the observation of increased levels of unsaturated fatty acids in seeds from that region. We also constructed a gene co-expression network to investigate regional variation in network modules. The results of this analysis revealed regional differentiation for eight of twelve modules, but no clear relationship with oil biosynthesis. Overall, the differential expression of *SAD* genes offers a partial explanation for the observed differences in seed oil composition between Texas and Canada, while the expression patterns of other metabolic genes suggest complex regulation of fatty acid production and usage across latitudes.

## 1. Introduction

Clinal patterns of phenotypic variation are common in nature and often track environmental gradients such as latitude or elevation (De Frenne et al., 2013; Halbritter et al., 2018). Such clines are often attributed to local adaptation, particularly for traits closely associated with life history, due to the importance of matching life history transitions with the local environment (De Frenne et al., 2013; Donohue et al., 2010; Fabian et al., 2015; Stinchcombe et al., 2004). Given the likely importance of traits that co-vary with environmental clines, there is particular interest in dissecting their genetic basis. Such information can provide insight into the processes that influence the distribution of adaptive genetic variants across species ranges and tolerance to stressful conditions, especially in the context of climate change (Ahuja et al., 2010; Atkins and Travis, 2010).

A particularly intriguing example of clinal trait variation in plants relates to seed oil composition (Linder, 2000; Sanyal et al., 2018). Seed oils are composed of both saturated and unsaturated fatty acids (FAs), the relative proportions of which are thought to influence germination timing and the amount of energy available to seedlings across temperatures (Linder, 2000). While saturated FAs provide more energy than unsaturated FAs, the lower melting point of the unsaturated FAs increases energy availability at cooler temperatures. Notably, a negative relationship between latitude and the proportion of saturated FAs in seeds has been found for several taxa (Linder, 2000; Sanyal et al., 2018). Seed oils from tropical plant species near the equator tend to have significantly greater proportions of saturated FAs than temperate plants in higher latitudes (Linder, 2000; Manos and Stone, 2001). It has thus been suggested that variation in seed oil composition is an adaptation to the temperatures most often experienced during germination (Linder, 2000; Sanyal and Decocq, 2016; Sanyal and Linder, 2013).

One of the best-studied examples of clinal variation in seed oil composition is in wild sunflower (*Helianthus annuus* L.). Across their native range, wild sunflower populations vary latitudinally in the proportion of saturated FAs in their seed oil – i.e., southern populations tend to produce seeds with a greater proportion of saturated FAs than northern populations (Figure 1; Linder, 2000; McAssey et al., 2016). Given that this pattern is maintained in a common garden, it seemingly results from genetic differentiation across the species range. Evidence for a trade-off in germination timing at high and low temperatures between populations from the extreme ends of the gradient has also been observed, consistent with the idea that natural selection on oil composition (due to its effects on germination and/or seedling establishment) is responsible for producing this pattern (Linder, 2000).

**Figure 1:**
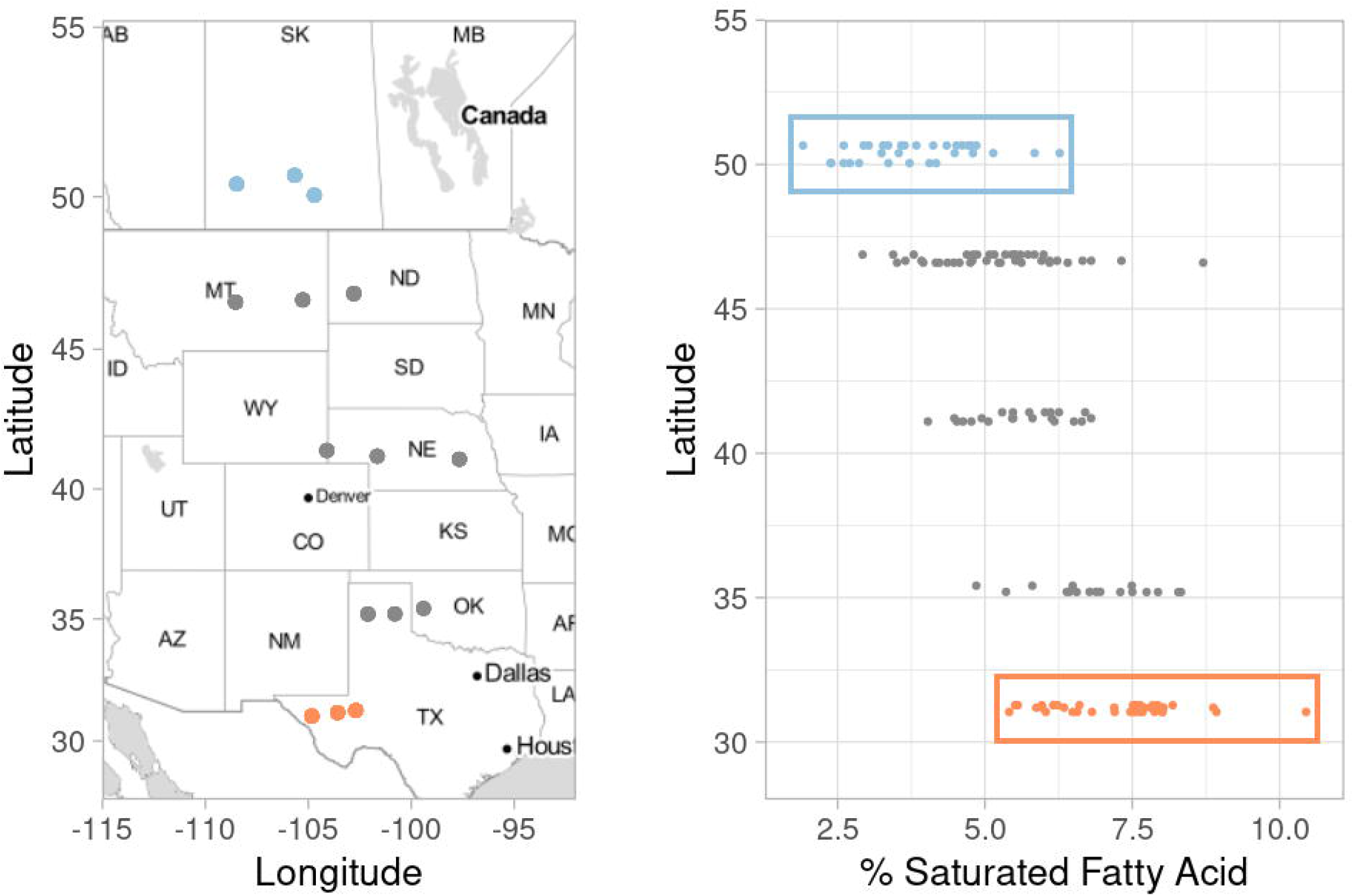
The proportion of saturated fatty acids in seed oils of wild sunflower varies along a latitudinal gradient. A: Populations of wild sunflowers included in the work of McAssey et al. (2016). Orange and blue dots represent the populations from Texas and Canada that were included in this study. Grey points represent populations studied by McAssey et al. (2016) but not included in this study. B: Concentration of saturated fatty acids in seed oils from wild sunflower populations across a latitudinal gradient. Samples from Texas are colored orange and grouped within an orange box while samples from Canada are colored blue and grouped within a blue box. The proportion of saturated fatty acids is significantly higher in seeds from Texas populations than from Canada populations as described in McAssey *et al*. (2016) and denoted by an asterisk.

In addition to being a model for ecological studies, wild sunflower is the progenitor of cultivated sunflower (also *H. annuus*), which is one of the world’s most important oilseed crops. Cultivated sunflower was initially domesticated over 4,000 years ago in what is now the east-central USA as a source of edible seeds and for cultural purposes (Heiser, 1978; Park and Burke, 2020; Rieseberg and Seiler, 1990), and was later subjected to intense selection for increased seed oil concentration, as well as a shift in FA composition (Fick and Miller, 1997). Sunflower oil is a naturally rich source of oleic and linoleic acids (18:1 and 18:2 respectively, where the numbers represent the length of the carbon chain and the number of double bonds present [i.e., the degree of FA unsaturation]). These two FAs combine to account for ca. 85-90% of the total seed oil content, with the remainder being largely made up of the fully saturated palmitic (16:0) and stearic (18:0) acids. In recent years, breeding efforts have increasingly focused on the development of high-oleic lines that produce seed oil composed of ≥ 80% oleic acid, which is valued for its health benefits and stability during storage (Miller et al., 1987).

Due to the ecological and economic importance of seed oil composition and the general importance of fatty acids in biological systems, the fatty acid metabolic pathway has been well-characterized (Bates et al., 2013). In plants, palmitic (16:0) and stearic (18:0) acid are the first long-chain FAs produced during FA synthesis. Stearoyl-ACP desaturases (SAD) then convert palmitic and stearic acid into monounsaturated palmitoleic (16:1) and oleic acid. Fatty acid desaturases (FAD) then convert oleic acid into linoleic acid and other polyunsaturated fatty acids. The role of the enzyme-producing *SAD* and *FAD* genes in controlling oil composition has been well established in many plant species (Belide et al., 2012; Dar et al., 2017; Fofana et al., 2006; Liu et al., 2002; Rajwade et al., 2014; Thambugala and Cloutier, 2014). In cultivated sunflower, the regulation of these genes is responsible for many commercially important seed oil phenotypes, including the aforementioned high-oleic acid phenotype, which is conditioned by a mutation resulting in the downregulation of the seed-specific *FAD2-1* gene (Hongtrakul et al., 1998; Miller et al., 1987; Schuppert et al., 2006). Similarly, increased stearic acid content has been mapped to quantitative trait loci (QTL) containing *SAD* genes (Pérez-Vich et al., 2004; Pérez-vich et al., 2002). While regulation of *SAD* and *FAD* genes underlie these commercially important phenotypes in cultivated sunflower, less is known about how these and related genes contribute to observed variation in seed oil composition in wild populations.

As a follow-up to observations of latitudinal variation in seed oil composition among wild sunflower populations (McAssey et al. 2016; Figure 1), we sought to characterize transcriptomic variation in developing seeds of plants derived from the southern and northern extremes of the species range. Given what is known about the genetic basis of oil biosynthesis in cultivated sunflower, we asked: 1) which genes are differentially expressed between Texas and Canada and what are their putative biological roles?; 2) are genes related to oil metabolism, particularly members of the *SAD* and *FAD* gene families, differentially expressed between regions?; 3) are differentially expressed genes (DEGs) related to oil metabolism found within known oil QTL?; and 4) are DEGs related to oil metabolism found within co-expression network modules associated with geographic regions? To answer these questions, we sequenced the transcriptomes of developing seeds from the Texan and Canadian populations that were produced in the common garden experiment described by McAssey et al. (2016; Figure 1). Overall, several genes related to oil metabolism are differentially expressed across the range of wild sunflower. We found that *SAD* genes specifically are expressed at a significantly higher level in developing seeds from Canadian populations when compared to those from Texan populations, consistent with our expectations given the observed higher levels of desaturation at more northern latitudes. Additionally, many of the DEGs related to oil metabolism were found within oil QTL. Contrary to expectations, however, DEGs related to oil metabolism did not group into a single co-expression network module associated with region, which suggests oil metabolism genes do not act in a coordinated fashion to produce the observed seed oil phenotypes. Taken together, our results suggest that the regulation of seed oil metabolism across latitudes is a complex process that is influenced by many different genes.

## 2. Materials and Methods

### 2.1 Plant Materials

Seeds used in this study were produced via intrapopulation crosses in a common greenhouse environment during a previous experiment (McAssey et al., 2016). Briefly, seeds from six populations representing the southern and northern ends of the range of wild sunflower (three from Texas, USA and three from Saskatchewan, Canada; Figure 1, Supplemental Table 1) were germinated, allowed to establish, and then transplanted into pots in the greenhouse where they were arranged in a randomized fashion and grown to flowering. Flower heads were bagged once buds began to develop to prevent unintended cross-pollination. Because wild sunflowers are self-incompatible, pairs of individuals originating from the same population were manually cross-pollinated. This involved removing the pollination bags from the two focal plants, collecting pollen from both plants directly into a petri dish, and then using a paintbrush to apply this pollen to receptive florets of the same two plants. Because florets within an inflorescence open over time, pollination was performed over a period of days for each individual. Following pollination, heads were re-bagged to prevent contamination and plants were allowed to set seed. Fifteen days after the final pollination, eight developing achenes (i.e., single-seeded fruits) were collected from the center of each head, placed into 1.5 mL tubes, and frozen in liquid nitrogen. These tubes were then stored at −80C until RNA extraction.

### 2.2 RNA Extraction and Library Construction

The eight frozen achenes from each maternal plant were pooled into a single sample and ground with a mortar and pestle using liquid nitrogen and a small amount of polyvinylpolypyrrolidone (PVPP). The ground tissue was then transferred into a tube and placed in liquid nitrogen to keep it frozen. After removing individual samples from the liquid nitrogen for processing, 1 mL of Trizol was added to each tube. The contents were then mixed and allowed to incubate at room temperature for five minutes. Chloroform (300 μl) was then added to each sample and the tubes were manually shaken and then centrifuged at 12,000 G for 10 minutes. The aqueous phase of each sample was then removed via pipetting and transferred to a new tube. After mixing with a 0.53X volume of 100% ethanol, RNA was purified by processing the solution following the Qiagen RNeasy Plant Mini Kit (Valencia, CA, USA) protocol with on-column DNase digestion.

RNA quality was assessed using an Agilent Bioanalyzer (Santa Clara, CA). All samples (nine from Texas, eleven from Canada; Table 4.1) used for library construction had RIN values ≥ 8.5 out of 10. Libraries were constructed using a Kapa mRNA-seq kit (Kapa Biosystems, Wilmington, MA). This kit performs size selection using magnetic beads. Libraries were constructed to include a size range of approximately 200-500 bp, and size ranges were checked using a fragment analyzer (Advanced Analytical Technologies, Ankeny, IA). Individual libraries were then quantified via qPCR using Illumina (San Diego, CA) standards and equimolar amounts of each library were pooled into a single tube and submitted for Illumina NextSeq 500 SE75 sequencing at the Georgia Genomics and Bioinformatics Core (http://dna.uga.edu; Athens, GA). The resulting sequence data are available via the Short Read Archive (SRA) under BioProject PRJNA706177.

The resulting RNAseq data were processed using a custom bioinformatics pipeline (https://github.com/EDitt/Sunflower_RNAseq). Adapter sequences were first trimmed using Trimmomatic v0.36 with default settings (Bolger et al., 2014). Next, because, samples were split across 4 lanes during sequencing, reads from each lane were separately mapped to the XRQv1 sunflower genome assembly (Badouin et al., 2017) using the two-pass mapping method implemented in STAR v2.6.1c (Dobin et al., 2013). The program RSEM v1.3.1 (Li and Dewey, 2011) was then used to calculate expression levels for each gene in each sample prior to downstream analysis.

### 2.3 Differential expression analyses

DEGs were identified using edgeR v3.24.3 (Robinson et al., 2010). Only genes that were expressed at a threshold of at least one count-per-million in two or more samples per region were retained for this analysis. The model matrix used to estimate dispersions incorporated source population and region as factors. In addition, we tested for differential expression between population pairs within Texas and Canada, with the exception of TEX2, which only had one sample. The false discovery rate was controlled for by applying a Benjamini-Hochberg correction (Benjamini and Hochberg, 1995) to the set of *P*-values calculated by edgeR.

Gene ontology (GO) terms for each of the sunflower genes were obtained from the blast2go (Conesa and Götz, 2008) output available on the XRQv1 genome portal (https://www.heliagene.org/). GOseq v1.34.1 (Young et al., 2010) was then used to calculate enrichment of GO terms within the set of differentially expressed genes while accounting for transcript length bias and calculating *P*-values using Wallenius’ noncentral hypergeometric distribution (Wallenius, 1963) after which the false discovery rate was controlled for by applying a Benjamini-Hochberg correction to the raw *P*-values (Benjamini and Hochberg, 1995). Genes in the sunflower genome were also mapped to metabolic pathways using the Mercator4 tool (Schwacke et al., 2019) and DEGs mapping to the X4 Lipid Metabolism basic process were arranged and displayed in a heatmap using the heatmap.2 function from the R package gplots v3.1.1 (Galili, 2021).

### 2.4 QTL co-localization

We determined the locations of DEGs relative to the positions of previously mapped QTL related to fatty acid biosynthesis that were compiled in Badouin et al. (2017) from several other previously published works (Ebrahimi et al., 2008; Pérez-Vich et al., 2016; Premnath et al., 2016). The HanXRQv1 base pair coordinates of QTL from those three studies were compiled in Badouin et al. (2017) and used in this study. The genomic coordinates of oil QTL from Burke et al., 2005 were determined by: (1) identifying the simple-sequence repeat (SSR) markers surrounding each QTL, (2) locating those SSR markers on the consensus genetic map of the sunflower genome (Bowers et al., 2012), (3) identifying the nearest single-nucleotide polymorphism (SNP) to each of those markers, and (4) locating the target sequences for each SNP probe (Bachlava et al., 2012) on the sunflower XRQv1 genome via blast and determining their base pair coordinates. All genes within the intervals of interest were then extracted and cross-referenced with our list of DEGs (Supplemental Data Set 1).

### 2.5 Co-expression network analysis

We built a co-expression network using CemiTool v3.12 (Russo et al., 2018) in R to test for associations between geographic region (i.e., Texas vs. Canada) and individual co-expression modules. We generated a signed network using Pearson correlation coefficients. Invariant genes were filtered out from module construction using default settings and modules were then tested for their association with region. Hub genes were identified as the top 5 most connected genes within each module. Module expression plots were recreated in R by normalizing expression data to log2 transcripts-per-million (TPM), plotting each gene within the module, and averaging their expression values. Co-expression modules were then tested for enrichment of GO terms using CemiTool’s over-representation analysis.

## 3. Results and Discussion

### 3.1 Which genes are differentially expressed between Texas and Canada and what are their putative biological roles?

#### 3.1.1 Transcriptome sequencing and sample clustering

Sequencing of the 20 RNA libraries resulted in ca. 559M total reads, of which ~95% could be assigned to a sample. One sample with less than 1M reads was removed prior to the analysis. For the remaining libraries, there was an average of 26.4M reads per library, ranging from 9.3–85M reads (Supplemental Figure 1). Just over half (29,425 of 58,138; 50.6%) of the genes met the minimum expression threshold of ≥1 count-per-million in at least 2 samples and were retained for analysis. Multidimensional scaling (MDS) of samples based on gene expression was used to assess the level of dissimilarity between samples (Figure 2A). Samples from Texas were quite similar to each other, though they did not form a tight cluster. In contrast, samples from Canada were differentiated from those in Texas, but formed two distinct clusters with individuals from CAN1 appearing distinct from those of CAN2 and CAN3 (Supplemental Figure 2).

**Figure 2:**
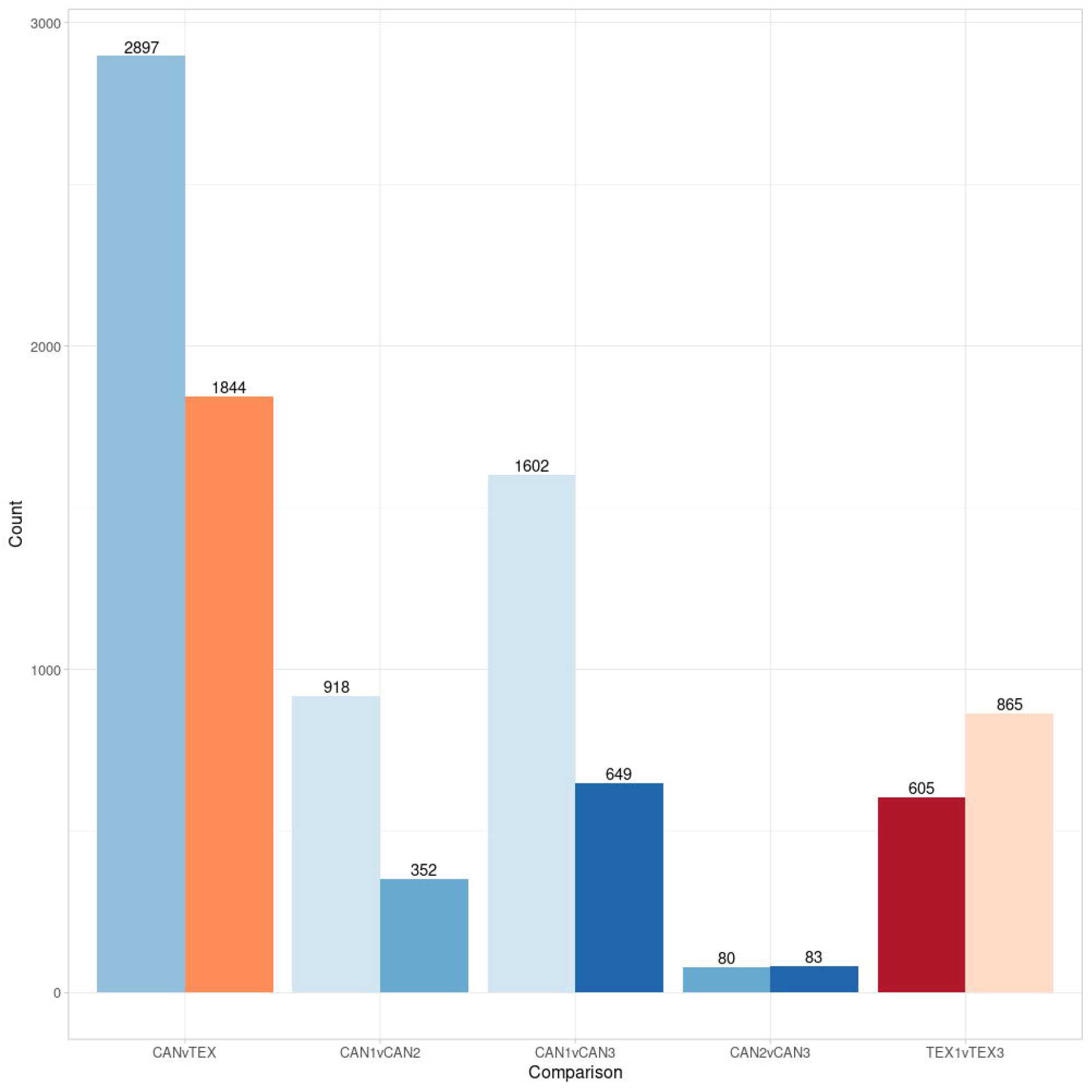
Barplot showing the number of differentially expressed genes between samples from Texas and Canada while also showing the number of differentially expressed genes between individual populations within those regions. Each pair of bars represents the total number of differentially expressed genes in each contrast. Individual bars represent the number of DEGs that were expressed more highly within that group in a given contrast. Shades of red represent populations from Texas while shades of blue represent populations from Canada.

A total of 4,741 genes were found to be differentially expressed between Texas and Canada, including 2,897 that were upregulated in Canada and 1,844 that were upregulated in Texas (Figure 2B, Supplemental Data Set 2). A smaller number of DEGs were identified in pairwise contrasts between populations within regions (Figure 2B). Consistent with the MDS plot, the CAN2 vs. CAN3 comparison had relatively few DEGs while the CAN1 vs. CAN2 and CAN1 vs. CAN3 comparisons had much larger sets of DEGs.

As these individuals were grown in a common garden, variation in gene expression is likely due to genetic differences between populations. The clear differences in gene expression between populations from Texas and Canada were expected based on evidence that wild sunflower exhibits north-south divergence in terms of both genetic and phenotypic variation (Blackman et al., 2011; McAssey et al., 2016; Park and Burke, 2020). These patterns likely result, at least in part, from selection for adaptation to the drastic environmental differences between Texas and Canada; such selection might also have contributed to variation in the expression of genes related to environmental adaptation (Akman et al., 2016; Lasky et al., 2014).

#### 3.1.2 GO term enrichment within DEGs

Among the 4,741 genes that were differentially expressed in samples from Texas vs. Canada, 16 GO terms were found to be significantly enriched (Supplemental Data Set 3, Table 1). Most of the enriched GO terms fell into the molecular function ontology category, with fewer falling into the biological process category, and only a single term being a part of the cellular component category. The enrichment of the single cellular component term, membrane (GO:0016020), could be related to differences in seed oil composition, as the fatty acid composition of membranes is critical for maintaining membrane fluidity at different temperatures (Falcone et al., 2004). However, GO terms directly related to fatty acid production, such as lipid metabolism (GO:0006629), omega-6 fatty acid desaturase activity (GO:0045485), stearoyl-ACP-desaturase activity (GO:0102786), and several others, were not found to be enriched within this set of DEGs, even prior to the multiple test correction.

**Table 1.** GO term enrichment among genes differentially expressed between Texas and Canada.

| Category | Term | Ontology | P-value | Number of DEGs In category | Total number of genes in category |
| --- | --- | --- | --- | --- | --- |
| GO:0000166 | Nucleotide binding | Molecular function | 5.47E-06 | 190 | 965 |
| GO:0043531 | ADP binding | Molecular function | 3.32E-05 | 48 | 167 |
| GO:0006952 | Defense response | Biological process | 7.87E-05 | 69 | 285 |
| GO:0016020 | Membrane | Cellular component | 7.87E-05 | 491 | 3010 |
| GO:0016310 | Phosphorylation | Biological process | 0.0001430701155 | 144 | 740 |
| GO:0016301 | Kinase activity | Molecular function | 0.0001504724832 | 120 | 590 |
| GO:0016740 | Transferase activity | Molecular function | 0.0111670116 | 136 | 723 |
| GO:0006468 | Protein phosphorylation | Biological process | 0.01324937678 | 155 | 878 |
| GO:0005342 | Organic acid transmembrane transporter activity | Molecular function | 0.02089174127 | 5 | 5 |
| GO:0004559 | Alpha-mannosidase activity | Molecular function | 0.02089174127 | 8 | 13 |
| GO:0006013 | Mannose metabolic process | Biological process | 0.02089174127 | 8 | 13 |
| GO:0005524 | ATP binding | Molecular function | 0.02089174127 | 357 | 2313 |
| GO:0004672 | Protein kinase activity | Molecular function | 0.02391560147 | 111 | 601 |
| GO:0003824 | Catalytic activity | Molecular function | 0.02778733722 | 64 | 302 |
| GO:0016787 | Hydrolase activity | Molecular function | 0.02840233826 | 138 | 764 |

### 3.2 Are genes related to oil metabolism, particularly members of the *SAD* and *FAD* gene families, differentially expressed between regions?

#### 3.2.1 Differential expression analysis of genes in the lipid metabolism pathway

While the *SAD* and *FAD* genes are the primary genes responsible for modulating the degree of fatty acid (de)saturation in plant seed oils (Bates et al., 2013), we also examined the expression of genes within the fatty acid metabolism pathway more broadly. A total of 850 genes in the sunflower XRQv1 genome were found with the Mercator4 X4 Lipid Metabolism pathway, 55 of which were differentially expressed between Texas and Canada based on our data. Interestingly, when samples were clustered based on the expression of these 55 genes, they did not cluster entirely by region. TX1-1 and TX1-4 clustered with the majority of CAN samples while CAN3-3 clustered with the rest of the TX samples (Figure 3). This could indicate that another factor, aside from location of origin, influences the observed expression of genes within this pathway. While care was taken to ensure that seeds from each individual were sampled at the same developmental stage, it is also possible that minor differences in seed maturity or temperature during development could be influencing the expression of these genes (Byfield and Upchurch, 2007; Fofana et al., 2006; Li et al., 2015; Wang et al., 2019).

**Figure 3:**
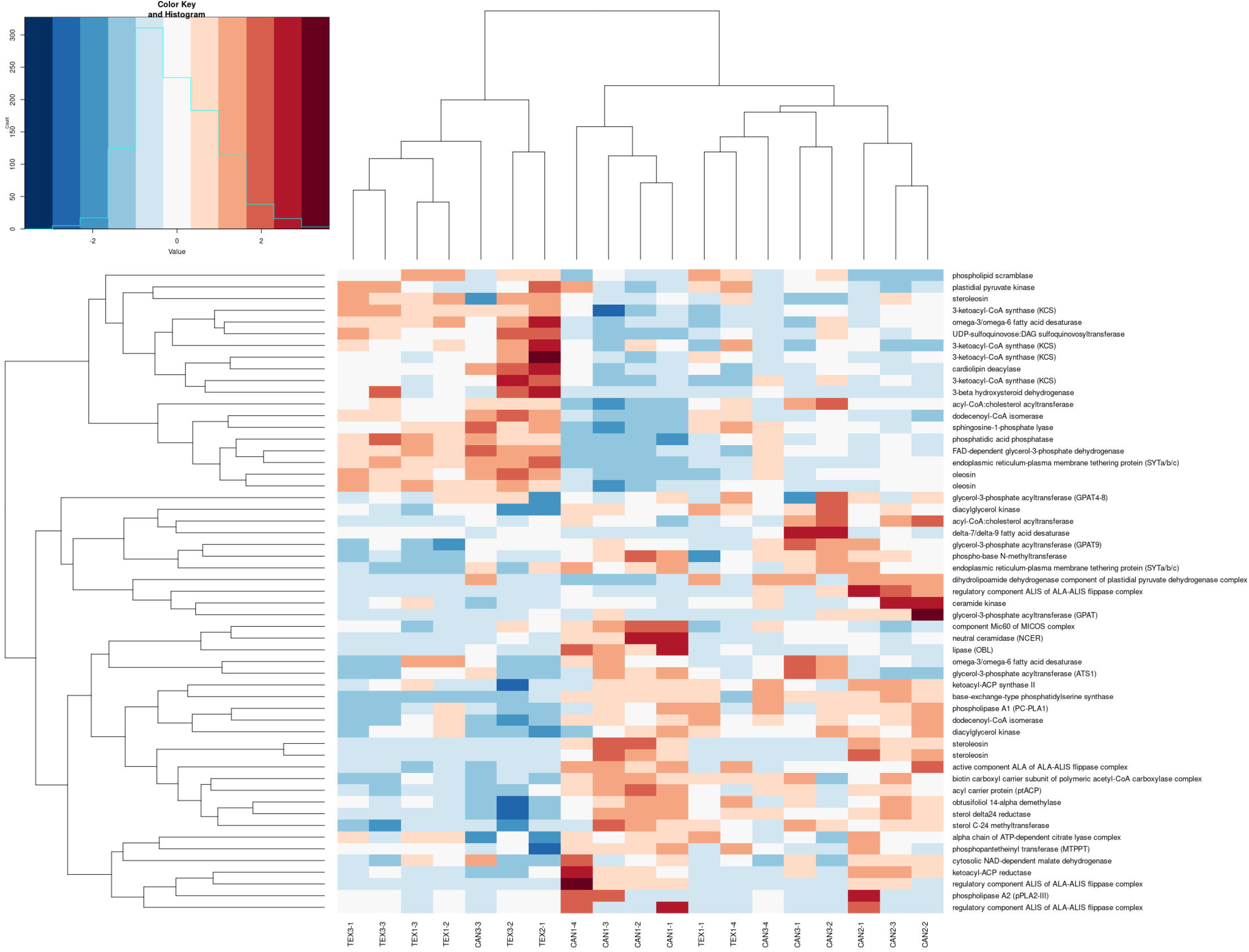
Heatmap of 55 DEGs that map to the Mercator4 X4 Lipid Metabolism pathway. Expression values are log transformed transcripts-per-million. Samples were clustered based on a dissimilarity matrix from the expression of these 55 genes.

Many of the DEGs within the lipid metabolism pathway play a role in fatty acid biosynthesis and utilization. For example, several 3-ketoacyl-CoA synthase (*KCS*) genes are more highly expressed in samples from Texas vs. Canada. *KCS* genes are involved in the elongation step during the synthesis of very-long-chain fatty acids (Gu et al., 2020a; Millar and Kunst, 1997). Drought stress is known to trigger production of these very-long-chain fatty acids, and while they don’t affect the overall degree of seed oil saturation, expression of these genes can alter the levels of individual fatty acids within seed oils (Gu et al., 2020a; Guo et al., 2020; Hashim et al., 1993). Members of the ALA-ALIS flippase complex are also differentially expressed and are involved in the translocation of phospholipids between membrane bilayers (Nintemann et al., 2019). A single delta-7/delta-9 fatty acid desaturase is seemingly more highly expressed in CAN3-1 and CAN3-2 than in other CAN samples, but otherwise does not vary substantially between Canada and Texas. Overall, the fact that several lipid metabolism genes are differentially expressed between regions indicates that there is complex regulation and usage of fatty acids involved in adaptation to the specific environmental conditions experienced in Texas vs. Canada. The ratio of unsaturated vs. saturated fatty acids in seed oil is likely only one part of a complex biological system.

#### 3.2.2 Expression patterns of *SAD* and *FAD* genes

The expression patterns of *SAD* and *FAD* genes in particular were examined to determine if they showed evidence of differential regulation consistent with the phenotypic differences between regions. Two *SAD* genes and 37 *FAD* genes were expressed in our developing seed samples. The two *SAD* genes and one *FAD* gene were highly expressed in these seed samples (Figure 4). Looking across the range, the two *SAD* genes were expressed at significantly higher levels in Canada vs. Texas (*P*-value = 0.026 and *P*-value = 0.027; Figure 4), although *P*-values were no longer significant after correcting for multiple comparisons in transcriptome-wide analyses. In contrast, the most highly expressed *FAD* gene, presumably the seed-specific oleic acid desaturase *FAD2-1,* was not differentially expressed between regions. While several of the more lowly expressed *FAD* genes did exhibit differential expression among regions (Supplemental Data Set 4), they were expressed at such low levels compared to *FAD2-1*(between 0 and 560 TPM, average 10.6 TPM) it is unlikely that they substantially impact seed oil composition.

**Figure 4:**
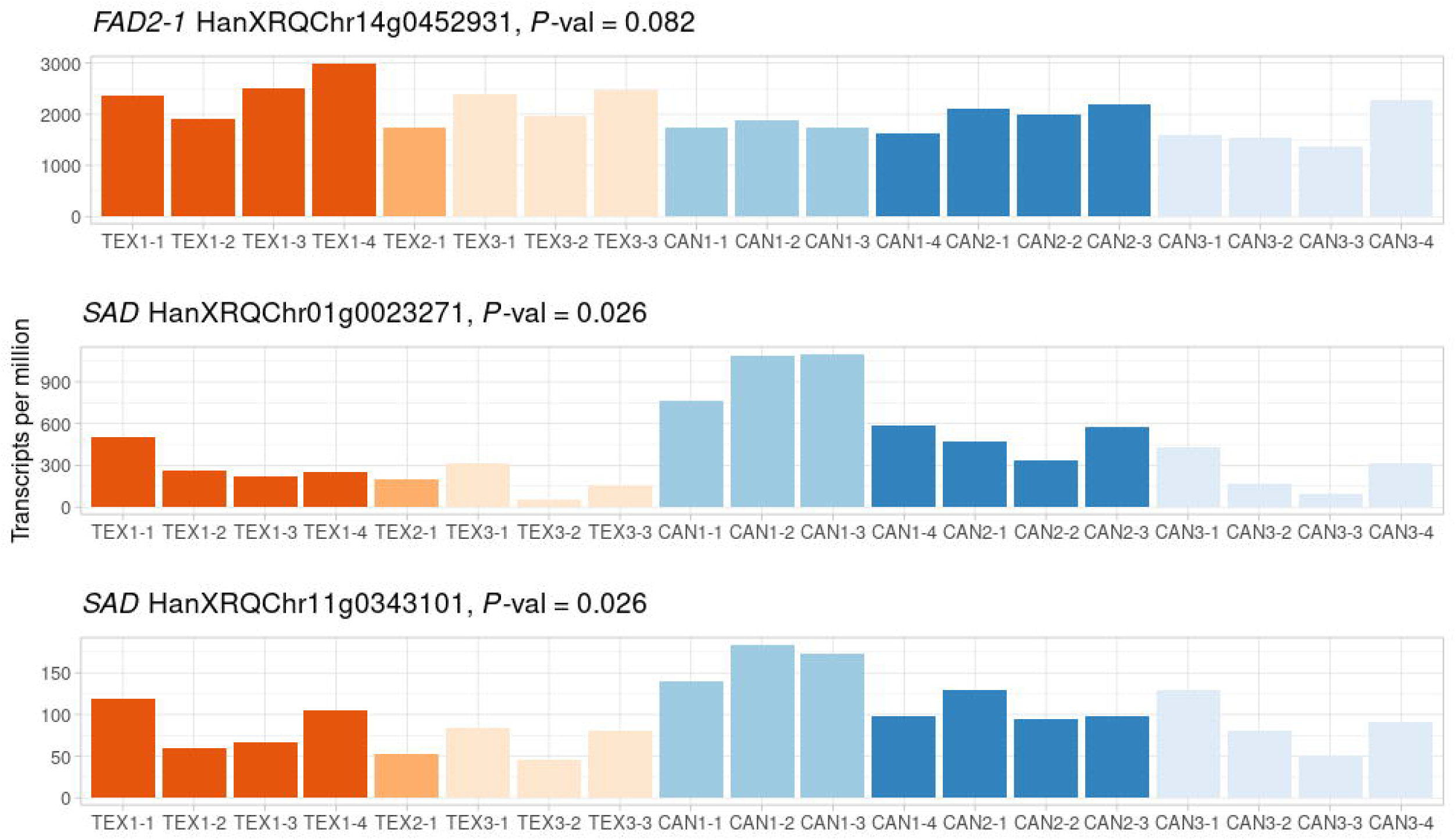
Comparison of the two *SAD* genes and the most highly expressed *FAD* gene (*FAD2-1*) among individuals from Texas and Canada. Gene expression is scaled to transcripts per million. *P*-values are uncorrected values obtained from the differential expression analysis.

Based on the functional role of *SAD* genes reported in other studies (e.g., Cantisán et al., 2000; Pérez-vich et al., 2002; Pleite et al., 2006; Salas et al., 2004), the potentially increased *SAD* gene expression in developing seeds from Canada compared to Texas is consistent with the observed phenotypic patterns showing higher unsaturated fatty acid content in seeds from Canadian populations (McAssey et al. 2016). Because the transition from fully saturated fatty acids to monounsaturated fatty acids is solely mediated by the SAD enzyme (Bates et al., 2013); it is thus reasonable to expect that increased expression of *SAD* genes would increase SAD enzyme activity, and convert more saturated fatty acids into monounsaturated fatty acids. While our data cannot definitively show that increased *SAD* gene expression in Canadian sunflowers is the cause of higher seed unsaturated fatty acid content, our results provide strong candidate genes for future investigation.

### 3.3 Are DEGs related to oil metabolism found within oil QTL?

#### 3.3.1 DEG colocalization with oil QTL

We also asked if DEGs were enriched within known sunflower oil QTL from previous mapping studies. We determined the genomic location of DEGs relative to 51 oil QTL from previously published mapping studies in cultivated sunflower and 8 oil QTL from a wild and cultivated sunflower cross (Badouin et al., 2017; Burke et al., 2005; Ebrahimi et al., 2008; Pérez-Vich et al., 2016; Premnath et al., 2016). There were 5,145 expressed genes located within oil QTL, 759 of which were DEGs. Among all oil QTL, DEGs were significantly under-enriched as compared to random expectation (−1.08-fold enrichment, 828.97 expected DEGs; *P*-value = 0.006), while within any individual oil QTL, DEGs were not significantly enriched.

We also asked whether any of the 55 differentially expressed oil metabolism genes colocalized with any oil QTL. Ten oil metabolism genes were found within oil QTL; three genes colocalized with the QTL for oleic acid content on chromosome 1 from a wild x cultivated cross, two genes colocalized with the QTL for oleic acid content on chromosome 3 from a wild x cultivated cross, and the remaining five genes individually colocalized with five additional QTL (Table 2). Two differentially expressed *FAD* genes are found within the oleic acid QTL on chromosome 1 (HanXRQChr01g0009721 and HanXRQChr01g0011241). While these genes act downstream of SAD and convert monounsaturated fatty acids to di- and tri-unsaturated fatty acids, it’s possible that their activity helps to increase the rate of conversion from fully saturated to monounsaturated fatty acids. Two differentially expressed *KCS* genes are also found within oil QTL; one within the oleic acid content QTL on chromosome 3 and the other on the palmitic acid content QTL on chromosome 8. As noted earlier, KCS genes contribute to the production and elongation of very-long chain fatty acids and are involved in abiotic stress tolerance (Gu et al., 2020b; Millar and Kunst, 1997). A differentially expressed ketoacyl-ACP synthase II (*KASII*) colocalizes with a palmitic, stearic, oleic, and linoleic acid content QTL on chromosome 9. *KASII* is a precursor in the production of palmitic and stearic acid and its activity may increase the content of saturated fatty acids. The *KCS* gene on chromosome 8 and *KAS* gene on chromosome 9 both colocalize with oil QTL derived from a cross between two cultivated sunflower lines. This result is consistent with the hypothesis that genes affecting seed oil characteristics vary in their expression within cultivated sunflower and may influence seed oil characteristics in wild sunflower, as well.

**Table 2:** Oil metabolism genes that colocalize with oil QTL. QTL coordinates are the range of base pairs on each chromosome across which the QTL can be found. QTL from cultivated x cultivated crosses were determined in Ebrahimi et al. (2008), Pérez-Vich et al. (2016), and Premnath et al. (2016) then compiled in Badouin et al. (2017). QTL from wild x cultivated crosses were determined in Burke et al. (2005).

| Gene | Annotation | QTL coordinates (base pair positions) | QTL phenotype (seed oil content) | QTL cross |
| --- | --- | --- | --- | --- |
| HanXRQChr01g0003891 | phospholipase A1 (PC-PLA1) | Chr01<br>125301302 - 135301303 | Oleic acid | Wild x cultivated |
| HanXRQChr01g0009721 | omega-3/omega-6 fatty acid desaturase | Chr01<br>125301302 - 135301303 | Oleic acid | Wild x cultivated |
| HanXRQChr01g0011241 | omega-3/omega-6 fatty acid desaturase | Chr01<br>125301302 - 135301303 | Oleic acid | Wild x cultivated |
| HanXRQChr01g0023271 | stearoyl-ACP desaturase | Chr01<br>125301302 - 135301303 | Linoleic, oleic, and stearic acid | Cultivated x cultivated |
| HanXRQChr03g0092231 | dodecenoyl-CoA isomerase | Chr03<br>129963181 - 168095999 | Oleic acid | Wild x cultivated |
| HanXRQChr03g0092661 | 3-ketoacyl-CoA synthase (KCS) | Chr03<br>129963181 - 168095999 | Oleic acid | Wild x cultivated |
| HanXRQChr06g0172791 | glycerol-3-phosphate acyltransferase (ATS1) | Chr06 6870150 - 29342604 | Palmitic, oleic, and linoleic acid | Wild x cultivated |
| HanXRQChr08g0228321 | 3-ketoacyl-CoA synthase (KCS) | Chr08 89935992 - 99935993 | Palmitic acid | Cultivated x cultivated |
| HanXRQChr09g0237851 | ketoacyl-ACP synthase II (KASII) | Chr09 0 - 6460295 | Palmitic, stearic, oleic, and linoleic acid | Cultivated x cultivated |
| HanXRQChr14g0445471 | acyl carrier protein (ptACP) | Chr14<br>123401162 - 133401163 | Oleic and stearic acid | Cultivated x cultivated |
| HanXRQChr14g0452931 | *FAD2-1* fatty acid desaturase | Chr14<br>140585231 - 158557967 | Tocopherol and oleic acid | Cultivated x cultivated |
| HanXRQChr17g0540371 | obtusifoliol 14-alpha demethylase | Chr17 17637974 - 25120357 | Palmitic acid | Wild x cultivated |

### 3.4 Are DEGs related to oil metabolism found within co-expression network modules associated with geographic regions?

#### 3.4.1 Gene co-expression network analysis

To further examine how gene expression varies between Canada and Texas, we built a gene coexpression network and tested for module-specific associations with geographic regions. The variance filtering step reduced the number of genes used for module construction to 3,115. These genes clustered into 12 co-expression modules, 8 of which exhibited significant differences in expression between regions (Table 3, Supplemental Data Set 5). We then conducted a GO term enrichment analysis on each module and found that no modules were significantly enriched for GO terms related to oil metabolism, although module M6 was suggestive of enrichment for genes within the GO category GO:0006633 lipid metabolic process (*P*-value = 0.06, Supplemental Data Set 5). This module did not, however, exhibit significant region-specific expression.

**Table 3:** Module information from the construction of a gene co-expression network.

| Module | # of genes in module | *P*-value association with Texas | *P*-value association with Canada | # of genes overlapping with oil QTL | Enrichment score |
| --- | --- | --- | --- | --- | --- |
| M1 | 797 | 0.0006 | 0.0006 | 108 | 2.79 |
| M2 | 530 | 0.0006 | 0.0006 | 84 | 2.86 |
| M3 | 489 | 0.1667 | 0.4444 | 56 | -2.16 |
| M4 | 242 | 0.7626 | 0.8604 | 47 | -0.94 |
| M5 | 198 | 0.0110 | 0.0114 | 38 | -4.99 |
| M6 | 177 | 0.9935 | 0.9947 | 29 | 0.62 |
| M7 | 169 | 0.0110 | 0.0110 | 27 | -1.42 |
| M8 | 126 | 0.0076 | 0.0073 | 25 | -4.02 |
| M9 | 118 | 0.0076 | 0.0073 | 22 | -3.43 |
| M10 | 106 | 0.6740 | 0.6738 | 16 | 0.96 |
| M11 | 85 | 0.0110 | 0.0114 | 25 | 1.52 |
| M12 | 78 | 0.0298 | 0.0425 | 15 | -1.39 |

The five genes with the highest connectivity within each module were identified as ‘hub’ genes (Supplemental Data Set 5). Many of these hub genes were of unknown function while very few were related to lipid metabolism. Module M7 included two GDSL esterase lipase genes and a non-specific lipid transfer gene as hub genes, while no other module had any hub genes that were seemingly related to lipid metabolism. We then looked for differentially expressed oil metabolism genes within each module, similar to our analysis of oil metabolism genes within QTL. Of the 55 oil-related DEGs, 11 were found within co-expression modules (Table 4). In addition, one of the *SAD* genes was also found in a co-expression module. Only the *FAD* gene HanXRQChr01g0011241 was found both within a co-expression module and anoil QTL, but this *FAD* gene was lowly expressed among all our samples (between 0.5 and 39.1 TPM, average 14.8 TPM).

**Table 4:** Differentially expressed oil metabolism genes found within co-expression network modules.

| Module | Gene | Annotation |
| --- | --- | --- |
| M1 | HanXRQChr05g0138441 | cytosolic NAD-dependent malate dehydrogenase |
| M1 | HanXRQChr05g0159681 | acyl-CoA:cholesterol acyltransferase |
| M1 | HanXRQChr06g0174791 | steroleosin |
| M1 | HanXRQChr06g0179691 | phospho-base N-methyltransferase |
| M1 | HanXRQChr14g0427531 | neutral ceramidase (NCER) |
| M3 | HanXRQChr01g0011241 | omega-3/omega-6 fatty acid desaturase |
| M3 | HanXRQChr08g0209891 | sphingosine-1-phosphate lyase |
| M3 | HanXRQChr17g0549281 | oleosin |
| M6 | HanXRQChr14g0458271 | dodecenoyl-CoA isomerase |
| M6 | HanXRQChr01g0023271 | stearoyl-ACP desaturase |
| M8 | HanXRQChr05g0131921 | alpha chain of ATP-dependent citrate lyase complex |
| M8 | HanXRQChr06g0172791 | glycerol-3-phosphate acyltransferase (ATS1) |

While genes related to oil metabolism were found within co-expression modules, we did not find large groups of oil metabolism genes that were co-expressed and also varied between regions. This suggests that the variation in seed oil composition between regions is not the result of coordinated gene regulation within a co-expression module. Instead, variation in seed oil composition is likely the result of several independent processes that do not necessarily act in a coordinated manner.

## Conclusions

Taken together, our results suggest that the observed differences in seed oil composition between regions likely result from the complex regulation of numerous genes involved in FA metabolism. Some evidence suggests that the expression of *SAD* genes in particular could play a role in altering seed oil composition and contribute to observed patterns of latitudinal variation in oil composition. Furthermore, several other genes related to lipid metabolism are differentially expressed across the range and can be found within oil QTL. While these genes could influence the FA composition of seeds, they could also be involved in other important biological processes. Overall, our results provide insights into the possible role of transcriptomic variation of many genes involved in oil metabolism, and specifically *SAD* and *FAD* genes, in shaping variation in seed oil composition across the range of wild sunflower while also providing avenues for future studies of this ecologically and agronomically important trait.

## Supporting information

Supplemental Data Set 1

Supplemental Data Set 2

Supplemental Data Set 3

Supplemental Data Set 4

Supplemental Data Set 5

## Acknowledgements

We would like to thank members of the Burke lab for critical reading of the manuscript, staff at the UGA Plant Biology Greenhouses for aiding with the growth of our plants, and the Georgia Genomics and Bioinformatics Core for sequencing our RNA libraries. This work was supported by a grant from the NSF Plant Genome Research Program (IOS-1444522) to JMB.

## Author Contributions

M.H.B. analyzed the data and wrote the paper. E.V.M grew the plants, extracted RNA, prepared the sequencing libraries, and contributed to writing the paper. E.L.D. assisted with analyses and contributed to writing the paper. J.M.B. contributed to writing the paper and provided support and funds for the project.

