## Supplemental Information for "Transcriptomics of developing wild sunflower seeds from the extreme ends of a latitudinal gradient differing in seed oil composition"

**Supplementary Table 1:** Accession numbers, location data, and number of plant samples used in this study.

| Sample ID | USDA PI# | Location | Latitude | Longitude | N |
|---|---|---|---|---|---|
| TX1 | 413160 | Texas, USA | 31.03972222 | -104.8302778 | 4 |
| TX2 | 664692 | Texas, USA | 31.18916667 | -103.5780556 | 1 |
| TX3 | 468476 | Texas, USA | 31.27277778 | -102.6922222 | 4 |
| CAN1 | 592311 | Saskatchewan, Canada | 50.39361111 | -108.4802778 | 4 |
| CAN2 | 592316 | Saskatchewan, Canada | 50.66 | -105.6647222 | 3 |
| CAN3 | 592320 | Saskatchewan, Canada | 50.0475 | -104.7072222 | 4 |

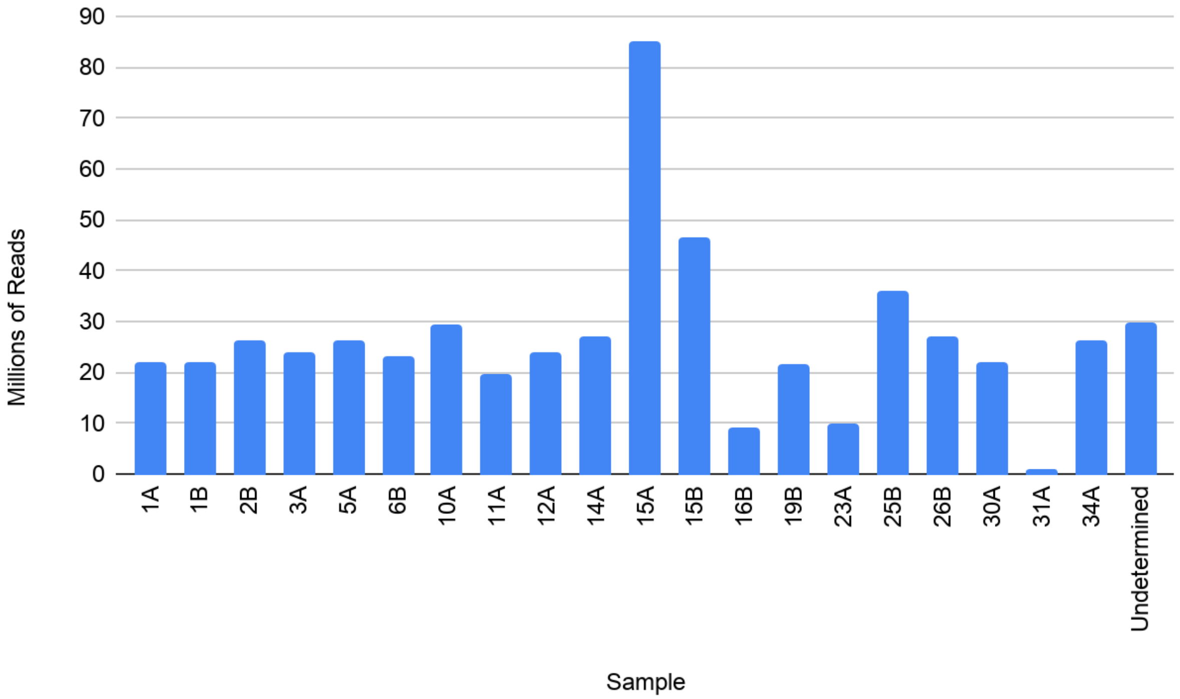


**Supplementary Figure 1:** Number of reads generated per sample from Illumina NextSeq run.

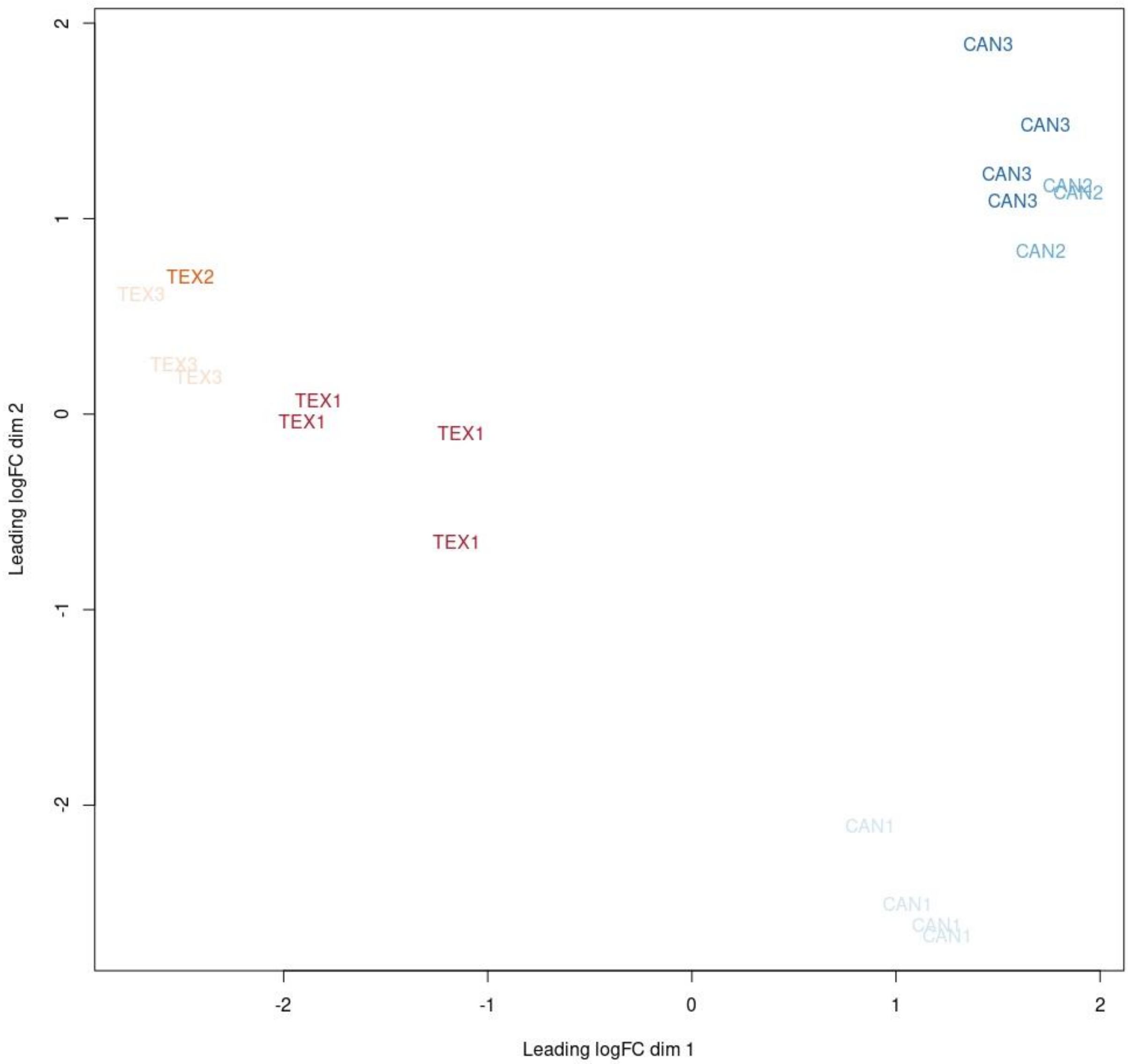


**Supplemental Figure 2:** Multidimensional scaling (MDS) plot generated from transcriptome data showing genetic differentiation between samples.
