## Supplemental Data Set 5 for "Transcriptomics of developing wild sunflower seeds from the extreme ends of a latitudinal gradient differing in seed oil composition": SupplementalDS5.html

CEMiTool


Code 

- Show All Code
- Hide All Code

### CEMiTool

### Report

#### Modules

#### Profile Plot

#### Gene Set Enrichment Analysis

#### Over Representation Analysis

### M1

### M2

### M3

### M4

### M5

### M6

### M7

### M8

### M9

### M10

### M11

### M12

#### Interaction Network

##### Please add interactions to the CEMiTool object

#### Parameters
